# Antisense oligonucleotide-based treatment of retinitis pigmentosa caused by mutations in *USH2A* exon 13

**DOI:** 10.1101/2020.10.06.320499

**Authors:** Ralph Slijkerman, Hester van Diepen, Silvia Albert, Margo Dona, Hanka Venselaar, Jingjing Zang, Stephan Neuhauss, Theo Peters, Sanne Broekman, Ronald Pennings, Hannie Kremer, Peter Adamson, Erik de Vrieze, Erwin van Wijk

**Affiliations:** MeiraGTx, 34-38 Provost Street, London, United Kingdom; Department of Otorhinolaryngology, Radboud University Medical Center, 6525 GA Nijmegen, the Netherlands; Human Genetics, Donders Institute for Brain, Cognition and Behaviour, Radboud University Medical Center, 6525 GA Nijmegen, the Netherlands; UCL, Institute of Ophthalmology, 11-43 Bath Street, London EC1V 9EL, United Kingdom; University of Zürich, Institute of Molecular Life Sciences, Zürich, Switzerland; Center for Molecular and Biomolecular Informatics, Radboud University Medical Center, 6525 GA Nijmegen, the Netherlands

**Author notes:** These authors contributed equally to this work (Ralph Slijkerman, Hester van Diepen). These authors contributed equally to this work (Peter Adamson, Erik de Vrieze, Erwin van Wijk).

**Keywords:** exon-skipping, QR-421a, retinitis pigmentosa, *USH2A*

## Abstract

Mutations in *USH2A*, encoding usherin, are the most common cause of syndromic and non-syndromic retinitis pigmentosa (RP). The two founder mutations in exon 13 (c.2299delG and c.2276G>T) collectively account for ^~^34% of *USH2A*-associated RP cases. Skipping of exon 13 from the *USH2A* transcript during pre-mRNA splicing presents a potential treatment modality in which the resulting transcript is predicted to encode a slightly shortened usherin protein. Morpholino-induced skipping of *ush2a* exon 13 in larvae of the previously published *ush2a* exon 13 zebrafish mutant resulted in the production of usherinΔexon13 and completely restored retinal function. RNA antisense oligonucleotides were investigated for their potential to specifically induce human *USH2A* exon 13 skipping. Lead candidate QR-421a induced dose-dependent exon 13 skipping in iPSC-derived photoreceptor precursors from a patient homozygous for the *USH2A* c.2299delG mutation. Intravitreal delivery of QR-421a in non-human primates showed that QR-421a penetrates the retinal outer nuclear layer and induces detectable levels of exon 13 skipping until at least 3 months post injection. In conclusion, QR-421a-induced exon skipping proves to be a highly promising treatment for RP caused by mutations in exon 13 of the *USH2A* gene.

## Introduction

Retinitis pigmentosa (RP) is a genetically and clinically heterogeneous disorder characterized by a progressive loss of visual function caused by the degeneration of the light-sensitive photoreceptor cells in the retina (Hartong *et al*, 2006). Although being designated as an orphan disease with an overall prevalence of 1:4,000 individuals, RP is the most common type of inherited retinal dystrophy (IRD), affecting ~125,000 patients within the European Union and almost two million individuals worldwide.

To date, mutations in over 100 genes are known to cause non-syndromic or syndromic RP (https://sph.uth.edu/Retnet/). It is estimated that autosomal recessively inherited RP (arRP) accounts for up to 60% of all RP cases (McGee *et al*, 2010). Mutations in *USH2A* collectively account for 7-23% of arRP cases, which can either result in non-syndromic arRP or in Usher syndrome (combination of RP and hearing impairment) (Rivolta *et al*, 2000; McGee *et al*, 2010). The mutations in this gene are mostly private and evenly distributed throughout the gene, but three mutations are derived from a common ancestor and are therefore seen more frequently: c.2299delG, p.(Glu767fs); c.2276G>T, p.(Cys759Phe) and c.7595-2144A>G, p.(Lys2532Thrfs) (Pennings *et al*, 2004; Aller *et al*, 2010; Vaché *et al*, 2012; Slijkerman *et al*, 2016). The c.2299delG and c.2276G>T mutations represent respectively 27.8% and 7.1% of all pathogenic *USH2A* alleles, and both reside in exon 13 (Baux *et al*, 2014). The auditory phenotype of patients with Usher syndrome can be partially compensated by providing patients with hearing aids or cochlear implants (Hartel *et al*, 2017). However, currently no treatment options exist for the progressive loss of vision associated with mutations in *USH2A*.

The poor understanding of the physiological role(s) of the usherin protein in photoreceptor cells, and the pathophysiological mechanisms underlying *USH2A*-associated RP, hamper the development of treatment that interferes with the disease mechanisms. The recent approval of Luxturna (voretigene neparvovec), a gene augmentation therapy for the treatment of patients with *RPE65*-associated retinal dystrophy (Bennett *et al*, 2016; Russell *et al*, 2017), has led to a paradigmatic shift in therapeutic research on retinal diseases, and provides hope for many visually impaired individuals worldwide. However, the development of an *USH2A* gene augmentation therapy is severely hampered by the size of the usherin-encoding sequence (15,606 nucleotides), which largely exceeds the cargo capacity of the currently used viral vehicles for gene delivery. An antisense oligonucleotide (AON) approach with naked intravitreal delivery could overcome this limitation. *In vitro* experiments demonstrated that AONs can be used to correct the aberrant pre-mRNA splicing caused by the c.7595-2144A>G mutation in *USH2A*, which leads to the inclusion of a pseudoexon in the mature *USH2A* transcript (Slijkerman *et al*, 2016). The promise for clinical application of AON-based therapies for inherited retinal dystrophies is currently under investigation (Xue and MacLaren 2020). QR-110 is a candidate oligonucleotide designed to treat patients with *CEP290*-associated Leber’s Congenital Amaurosis (LCA10). Treatment of LCA10 patients with QR-110 resulted in no serious adverse events and had a beneficial effect on vision after a single intravitreal delivery (Cideciyan *et al*, 2019).

In this study we explored AON-induced exon skipping as a potential treatment modality for patients with RP caused by mutations in exon 13 of the *USH2A* gene. As this exon consists of a multiplier of three nucleotides, skipping the exon would not disturb the open reading frame and could result in the production of a slightly shortened protein with residual function. Using our previously characterized *ush2a^rmc1^* zebrafish model (Dona *et al*, 2018), we demonstrate that usherin proteins lacking the amino acids encoded by exon 13 have sufficient residual function to restore the previously published retinal dysfunction. Using (patient-derived) cellular models and non-human primates, we identified and validated antisense oligonucleotide QR-421a as candidate molecule for the exon-skipping therapy described above. Currently, QR-421a is in clinical development (Xue and MacLaren 2020)

## Results

### Formation of EGF-like fusion domain after targeted *USH2A* exon 13 skipping

Wild-type usherin is predicted to contain ten EGF-lam domains (http://smart.embl-heidelberg.de/). EGF-lam domains typically contain eight cysteine residues that pairwise interact by a covalent disulfide bond, necessary for protein folding and stability. These EGF-lam domains in usherin harbor multiple protein truncating mutations. Also, in patients with *USH2A*-associated RP, 22 out of the 80 cysteine residues in these EGF-lam domains have been found to be mutated (*USH2A* LOVD mutation database, http://www.lovd.nl/USH2A), five of which reside within exon 13. Unpaired cysteine residues contain a reactive free thiol group that can induce unwanted multimerization or crosslinking with other proteins. The in frame skipping of exon 13 is predicted to result in the fusion of parts of EGF-lam domains 4 and 8 into a functionally related EGF-like domain (**Fig. 1A**). EGF-like domains contain six cysteine residues that together create three disulfide bonds by cysteines 1+3, 2+4 and 5+8 in the fused EGF-like domain (Wouters *et al*, 2005). There are 16 amino acids between the fifth and sixth cysteine residue in the EGF-like 4-8 fusion domain, which is different from the canonical spacing between cysteine residue 5 and 6 within EGF-like domains, namely 8 amino acids. However, 3D homology modelling predicted normal disulphide bridge formation within the EGF-like 4-8 fusion domain (**Fig. 1B**). In conclusion, molecular modelling warrants exploring the effect of exon 13 skipping at the level of visual function.

**Figure 1:**
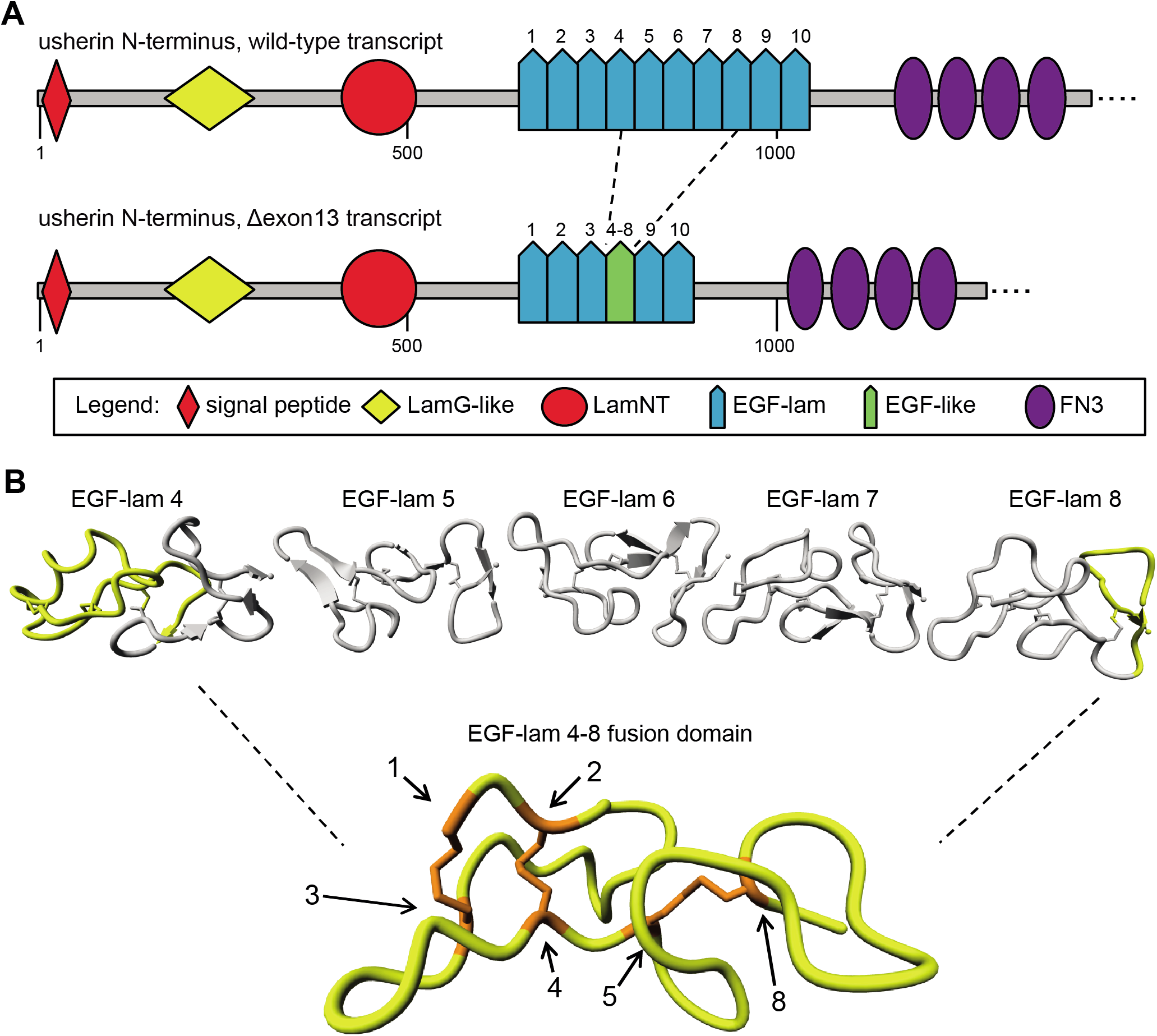
In silico modelling of usherin after exon 13 skipping. **(A)** Schematic representation of the domain architecture of wild-type usherin and usherinΔexon13. Individual EGF-lam domains are numbered. Skipping of exon 13 results in the exclusion of EGF-lam domains 5, 6 and 7 as well as the partial exclusion of EGF-lam domains 4 and 8. The remaining amino acids of EGF-lam domains 4 and 8 are predicted to form an EGF-like domain with six cysteine residues. The fusion site of this domain is located between the fifth cysteine residue of EGF-lam domain 4 and the eighth cysteine derived from EGF-lam 8. **(B)** 3D homology modelling predicts the formation of a stable EGF-like domain with normal disulphide bridge formations. The predicted structure of usherin EGF-lam domains 4 (left) and 8 (right) are shown. The amino acids that are encoded by USH2A exon 13 are depicted in grey and predicted to be absent after translation of *USH2A* Δexon13 transcripts. Cysteine residues that are present in the EGF-like fusion domain are numbered and indicated in orange. The cysteine residues numbered 1 to 5 are derived from EGF-lam domain 4, whereas residue 8 is derived from EGF-lam domain 8.

### AON-induced skipping of *ush2a* exon 13 in a mutant zebrafish model restores usherin protein expression and visual function

To validate exon 13 skipping as a potential therapeutic strategy, we employed our previously characterized *ush2a* zebrafish mutant (*ush2a^rmc1^*) that contains a frameshift-inducing mutation in exon 13 (Dona *et al*, 2018). We previously reported that electroretinogram (ERG) traces are significantly reduced in homozygous *ush2a^rmc1^* larvae, and that the usherin protein is absent from the retina, indicating that this mutant is a true null allele. The length of *USH2A* exon 13 is well conserved between human (642 nucleotides) and zebrafish (648 nucleotides) and the (spacing between) cysteine residues that are essential for EGF-lam domain formation is identical (**Fig. 2A**). Following the previously published guidelines for AON design (Aartsma-Rus, 2012; Slijkerman *et al*, 2018), six antisense phosphorodiamidate morpholino oligomers (PMOs) were designed to target the zebrafish *ush2a* exon 13 splice acceptor site, splice donor site, or exonic splice enhancer (ESE) motifs (**Fig. S1A**). The exon skipping potential of the PMOs was first investigated by injecting the individual PMOs in the yolk of 1- to 2-cell-stage *ush2a^rmc1^* embryos (**Fig. S1B**). Combined delivery of a low dose of two of the most potent PMOs, targeting different regions of *ush2a* exon 13, resulted in a more efficient skipping of *ush2a* exon 13 than observed for the individual PMOs (**Fig. S2A, S2B**). The combination of PMO1 and PMO2 appeared most potent after RT-PCR analysis, without causing severe toxic phenotypes, and was subsequently used to determine whether exon 13 skipping had an effect on the phenotypic outcome of the *ush2a^rmc1^* mutant (**Fig. 2B, Fig. S2C**).

**Figure 2:**
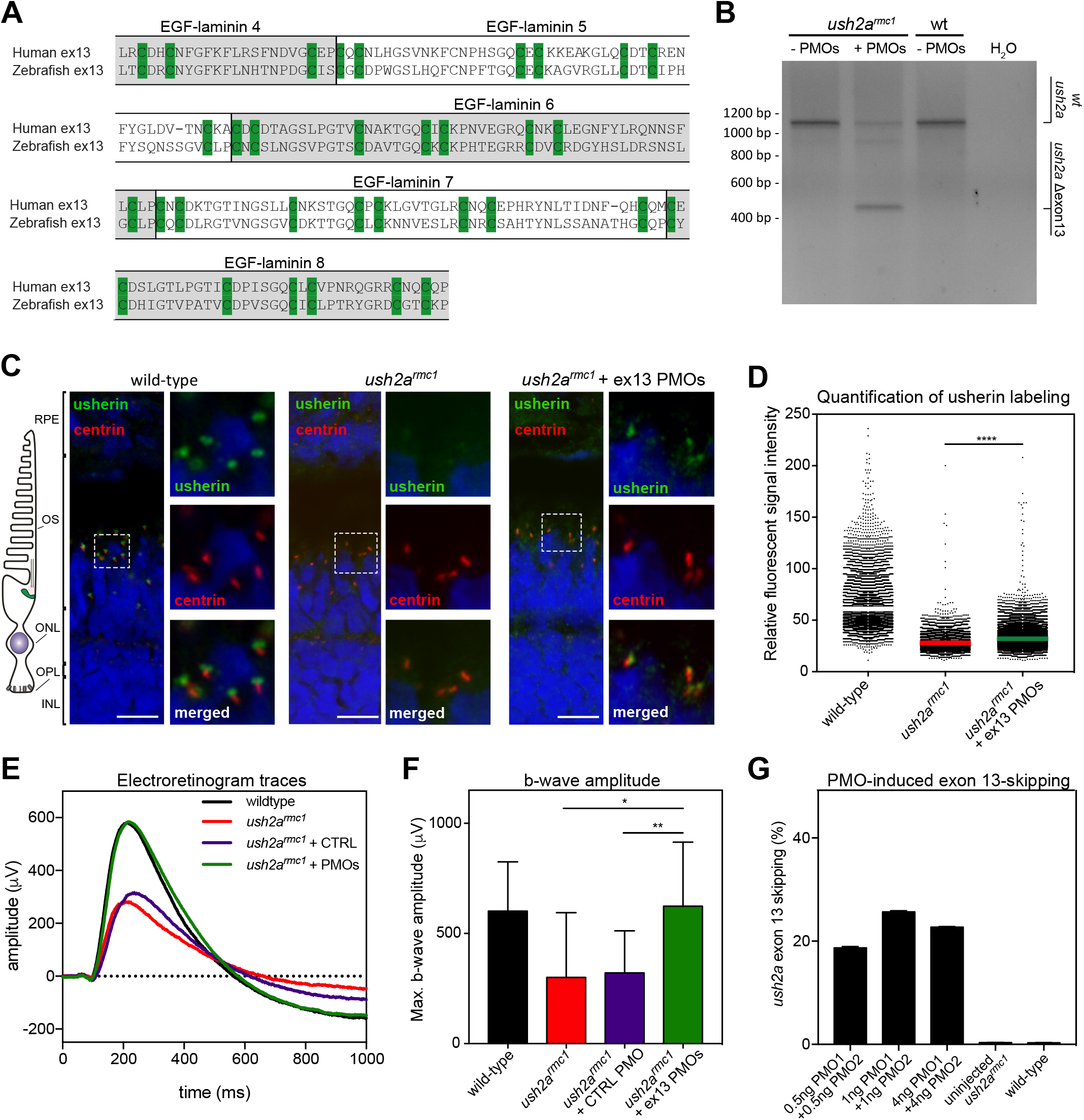
Morpholino antisense oligonucleotides mediate *ush2a* exon 13 skipping, usherinΔexon13 protein expression and restoration of ERG traces in a mutant zebrafish model. **(A)** Amino acid alignment of the sequences encoded by human and zebrafish *USH2A* exon 13. The (partial) EGF-lam domains are indicated. The cysteine residues required for 3D topology of the EGF-lam domains (green) are completely conserved between zebrafish and human. **(B)** Phosphorodiamidate morpholino oligonucleotides (PMO)-induced skipping of *ush2a* exon 13 in zebrafish larvae. *ush2a^rmc1^* mutant embryos were injected with a combination PMO1 and PMO2 (1pg of each). Investigation of *ush2a* pre-mRNA splicing at 3 dpf revealed the skipping of *ush2a* exon 13 upon injection of PMOs targeting *ush2a* exon 13. Uninjected and control PMO injected *ush2a^rmc1^* mutant zebrafish larvae and wildtype larvae were used as controls. **(C)** Subcellular localization of usherin in horizontal cryosections of larval (5 dpf) zebrafish retinae. Usherin was visualized with anti-usherin antibodies directed against the intracellular C-terminal tail of zebrafish usherin (green signal). Nuclei were stained with DAPI (blue signal), and the connecting cilium is labelled using anti-centrin antibodies (red). In wildtype larvae usherin is present at the photoreceptor periciliary membrane, adjacent to the connecting cilium. In homozygous *ush2a^rmc1^* larvae, no specific usherin signal could be detected. PMO-induced *ush2a* exon 13 skipping in *ush2a^rmc1^* mutant larvae resulted in partial restoration of usherin Δexon13 expression with the correct subcellular localization in the retina. **(D)** Scatterplot of the relative fluorescence intensity of anti-usherin staining in the periciliary region of all photoreceptors in the middle section of the larval zebrafish eye. The signal intensity is decreased in the *ush2a^rmc1^* retina compared to wild-types. Relative fluorescent signal intensity of anti-usherin staining is significantly increased in PMO-injected *ush2a^rmc1^* mutants as compared to uninjected mutants (**** P < 0.0001, Kruskal Wallis test followed by Dunns non-parametric post-test). **(E)** Average ERG b-wave traces from uninjected, control PMO injected, exon 13 PMO injected *ush2a^rmc1^* larvae, and wild-type controls at 5-6 dpf. PMO-induced skipping of *ush2a* exon 13 completely restored b-wave amplitudes in *ush2a^rmc1^* larvae as compared to uninjected or control PMO-injected mutants. **(F)** Normalized b-wave amplitudes recorded in uninjected or control PMO-injected *ush2a^rmc1^* larvae are significantly reduced as compared to ERG-traces from age- and strain-matched wild-type controls. Normalized b-wave amplitudes recorded in PMO-injected *ush2a^rmc1^* mutants are significantly improved as compared to ERG-traces from uninjected or control PMO-injected *ush2a^rmc1^* mutants, and do not significantly differ from wild-types (P > 0.99). Data is shown as mean ± SD, * P < 0.05, ** P <0.01, Kruskal Wallis test followed by Dunns non-paramentric post-test. **(G)** Quantification of ush2a Δexon13 transcripts in uninjected and PMO-injected zebrafish larvae at 3 dpf. Delivery of a higher dose of PMOs was not found to result in increased levels of *ush2a* Δexon13 transcripts. As the decrease of exon 13-containing transcripts was larger than anticipated based on the increase in *ush2a* Δexon13 transcripts (Fig. S2D), the amount of exon-skipping is presented relative to the amount of *ush2a* transcript levels in uninjected *ush2a^rmc1^* larvae. Abbreviations: OS: outer segment; ONL: outer nuclear layer; OPL: outer plexiform layer; IPL: inner plexiform layer; wt: wild-type; ush2armc1: zebrafish with exon 13 mutation; PMO: morpholino oligo; dpf: days post-fertilization; ERG: electroretinogram.

We first determined whether skipping of zebrafish *ush2a* exon 13 in homozygous *ush2a^rmc1^* mutant larvae resulted in the synthesis of a shortened usherin protein (usherinΔexon13). Antibodies directed against the intracellular region of zebrafish usherin were used to stain unfixed cryosections of wild-type larvae, uninjected *ush2a^rmc1^* larvae and *ush2a^rmc1^* larvae in which *ush2a* exon 13 skipping was induced by PMO injection (**Fig. 2C**, red signal). The photoreceptor connecting cilium was marked by antibodies directed against centrin (**Fig. 2C**, green signal). In wild-type larvae usherin localizes in the periciliary region, as expected. In the retina of uninjected *ush2a^rmc1^* larvae no usherin could be detected. In PMO-treated *ush2a^rmc1^* larvae a partial restoration of usherin expression was detected with a proper subcellular localization. The intensity of the anti-usherin fluorescence signals was quantified using an automated Fiji script. Skipping of *ush2a* exon 13 resulted in a small but statistically significant increase of the average fluorescence intensity in the periciliary region of photoreceptors as compared to untreated larvae from the same clutch (34.98 ± 0.14 (Δexon 13) versus 30.04 ± 0.16 (untreated); *p*<0.0001 (Kruskal Wallis and Dunns nonparametric test) (**Fig. 2D**). This corroborated that exon 13-skipping resulted in the synthesis of usherinΔexon13.

Next, we recorded electroretinograms (ERGs) from *ush2a^rmc1^* larvae that were treated with a combination of *ush2a* exon 13-targeting PMOs (n=25) or with a standard control PMO (n=14). Larvae from the same clutch were injected with a higher dose of PMO under the assumption that this would improve exon 13 skipping. As this increased the proportion of larvae with toxic phenotypes, ERGs were recorded from larvae that appeared visually normal in development. Uninjected age-, and strain-matched wild-types (n=10) and *ush2a^rmc1^* (n=11) larvae were used as controls. Uninjected and control PMO-injected *ush2a^rmc1^* mutant larvae demonstrated a significantly reduced b-wave amplitude as compared to age- and strain-matched wild-type larvae (*p*<0.05 (uninjected) and *p*<0.001 (control PMO-injected); Kruskal Wallis and Dunns nonparametric test) (**Fig. 2E, F**). PMO-induced skipping of *ush2a* exon 13 from *ush2a^rmc1^* larvae resulted in significantly increased b-wave amplitudes as compared to untreated or control PMO-injected *ush2a^rmc1^* larvae, which is indicative of an improved excitation of the secondary neurons. The ERG b-wave amplitudes recorded in *ush2a^rmc1^* larvae after injection with exon 13-targeting PMOs were not significantly different from those recorded in age- and strain-matched wild-type larvae (*p*>0.999) (**Fig. 2E, F**). Retrospective analysis of exon 13 skipping in larvae injected with low or high doses of PMOs revealed that increasing the PMO dose did not result in a significant gain in *ush2a* Δexon 13 transcripts, but rather decreased the number of full-length *ush2a* transcripts (**Fig. 2G, Fig. S2D**). At all tested doses of PMO, the levels of *ush2a* Δexon13 transcripts ranged between 18% and 26% of the amount of total *ush2a* transcripts observed in wildtype zebrafish. Together these data show that AON-induced skipping results in the formation and correct localization of an usherinΔexon13 protein with sufficient residual function to rescue visual dysfunction in *ush2a^rmc1^* zebrafish larvae.

### Identification of lead oligonucleotide QR-421a

Based on the ability of usherinΔexon13 to restore visual function in zebrafish, we aimed to develop antisense oligonucleotides (AONs) with the ability to induce skipping of exon 13 from human *USH2A* transcripts. A number of molecules were designed based on the bio-informatic analysis of the sequence of *USH2A* exon 13 and flanking intronic regions. Both the intron-exon boundaries, and the exonic splice enhancer (ESE) motifs within exon 13, identified using the SpliceAid webserver (Piva *et al*, 2009), were used as targets for AONs. Using *in silico* analysis, parameters for (lack of) secondary structure formation, thermodynamic properties, and sequence selectivity were taken into account to minimize potential off-target effects. The designed AONs were transfected in the retinoblastoma-derived WERI-Rb1 cell line (McFall *et al*, 1977) at a concentration of 200 nM and screened for their potential to induce *USH2A* exon 13 skipping (**Fig. S3**). As a result of these analyses, the best-performing 21-mer RNA antisense oligonucleotide sequence was selected. For further preclinical development, the molecule was synthesized as antisense RNA molecule with 2’-*O*-(2-methoxyethyl) sugar modification and a fully phosphorothioated backbone. This candidate was named QR-421a thereafter.

The target specificity of QR-421a was investigated after an *in silico* analysis. QR-421a shows no full complementarity to any mRNA, pre-mRNA or DNA target other than the anticipated region in *USH2A*. Partial complementarity to other genomic regions was only found with ≥2 mismatches. Only two off-target sequences were identified with 2 mismatches, residing in one intergenic and one intronic region, and are therefore not expected to influence gene splicing or expression. Other hits with >2 mismatches are not considered biological meaningful for a 21-mer splice modulation oligonucleotide, as a single mismatch in a similar AON was previously shown to already markedly decrease splice modulation efficiency (Garanto *et al*, 2019). Hence the risk for potential off-target effects due to the hybridization of QR-421a to targets other than the intended target, is considered negligible.

Next, WERI-Rb1 cells were treated with QR-421a gymnotically or with the aid of a transfection reagent to provide pharmacodynamic proof of concept for the *USH2A* exon 13 skipping potential using a transcript-specific quantitative ddPCR analysis. In both experiments, addition of QR-421a did not affect the total amount of (with and without exon 13) *USH2A* transcripts. Upon QR-421a transfection, a dose-dependent skipping of *USH2A* exon 13 was induced which was already evident at a concentration of 25 nM. An exon 13 skipping efficiency of ~60% was reached upon transfection of the highest concentration tested (200 nM) (**Fig. 3A**). After a gymnotic delivery of QR-421a in a concentration ranging from 10 to 50 μM, exon 13 skipping efficiencies ranging from 10 to 17% were observed (**Fig. 3B**). In both experiments, treatment of WERI-Rb1 cells with a control oligonucleotide did not induce skipping of exon 13, confirming that the observed exon skipping potential is specific for QR-421a (**Fig. 3A, B**). Amplification of *USH2A* exons 11 to 15 in QR-421a-treated WERI-Rb1 cells (200nM) revealed mainly transcripts lacking exon 13, which was confirmed by Sanger sequencing, but also two minor alternative products that were also identified in untreated WERI-Rb1 cells (**Fig. 3C, D**). One of these fragments lacked both exon 12 and 13, the other fragment contained only exons 11 and 15. Altogether these data show that QR-421a has the ability to enter proliferating WERI-Rb1 cells after transfection or even unaided, thereby inducing a concentration-dependent and specific skipping of *USH2A* exon 13 in the absence of inflammatory responses.

**Figure 3:**
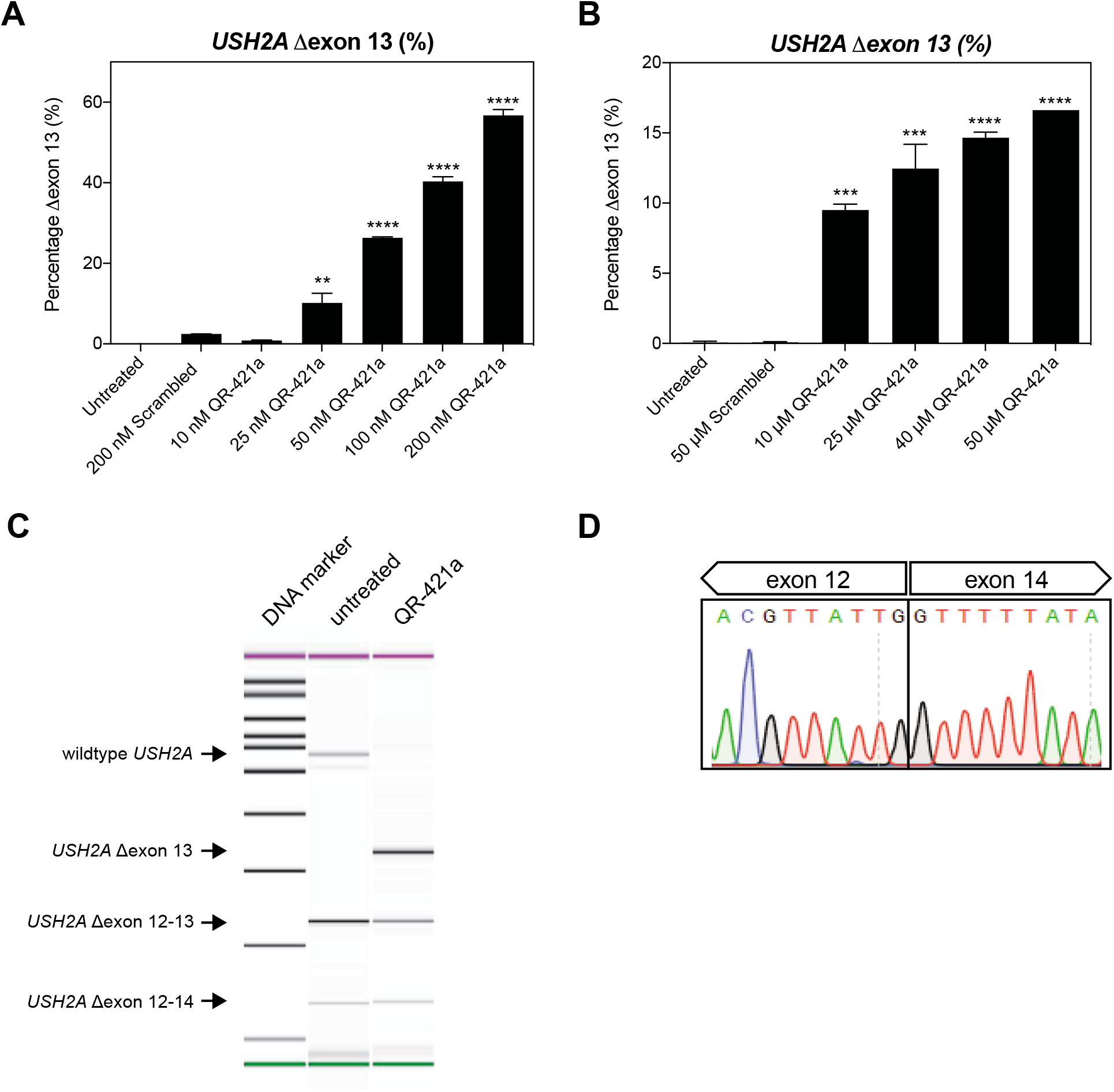
QR-421a shows a concentration-dependent increase of *USH2A* exon 13 skipping in WERI-Rb1 cells. WERI-Rb1 retinoblastoma cells were treated with different concentrations of QR-421a, using either **(A)** transfection or **(B)** gymnotic uptake. Untreated and scrambled control oligo treated cells were included as negative controls. Exon-skipping level was determined by quantification of *USH2A* Δexon13 and *USH2A* exon 13-containing transcripts using ddPCR. Treatment with QR-421a resulted in a significant concentration-dependent increase of *USH2A* Δexon13 transcripts. Data is shown as mean ± SD. Two biological replicates per treatment condition. Asterisks indicate significant differences with scrambled control oligo treated cells (*** P < 0.001; **** P<0.0001; One-Way ANOVA followed by Sidak’s Multiple Comparison Test). **C)** Representative image of amplicons present in untransfected and QR-421a transfected WERI-Rb1 cells. QR-421a is able to induce skipping of *USH2A* exon 13, and does not lead to the formation of alternatively spliced *USH2A* transcripts. Of note, *USH2A* Δexon 12-13 and Δexon 12-14 transcripts are already present in untreated WERI-Rb1 cells. **D)** Sanger sequencing traces of the *USH2A* Δexon 13 amplicon shown in **(C)** confirms that the sequence of exon 13 is lacking from the transcript.

### QR-421a treatment induces a concentration-dependent increase of *USH2A* exon 13 skipping in iPSC-derived photoreceptor progenitor cells

Photoreceptor progenitor cells (PPCs), differentiated from induced pluripotent stem cells (iPSCs) obtained from an *USH2A* patient with a homozygous c.2299delG mutation in exon 13, were used to assess the exon-skipping potential of QR-421a in a differentiated cell model with the appropriate genetic context. PPCs have been previously shown to be a valuable and clinically relevant tool for the evaluation of novel human-specific therapeutic strategies (Dulla *et al*, 2018; McDougald *et al*, 2016).

Initially, fibroblasts were reprogrammed into iPSCs and subsequently differentiated into PPCs. In order to validate that the cells had differentiated into PPCs, we assessed the expression levels of photoreceptor marker genes (*CRX*, *NRL*, *OPN1SW*, *OPN1LW* and *RHO*) by RT-qPCR analysis after 90 days of differentiation. As expected, the expression levels of photoreceptor marker genes were all significantly increased as compared to iPSCs, whereas the expression of the iPSC-specific marker gene *NANOG* was simultaneously decreased (**Fig. 4A**).

**Figure 4:**
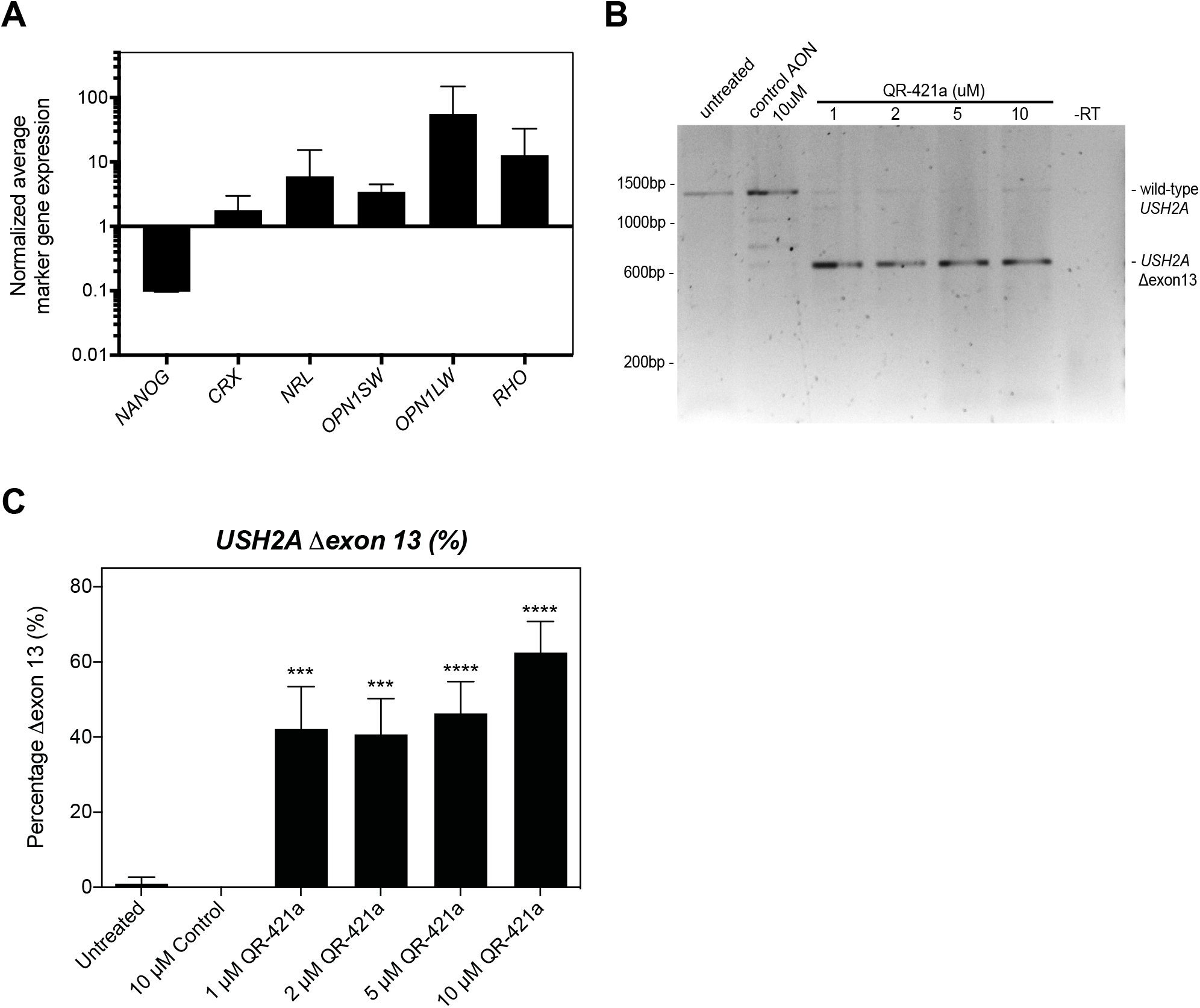
QR-421a treatment induces a dose-dependent increase of *USH2A* exon 13 skipping in iPSC-derived photoreceptor precursors cells (PPCs) from patients (*USH2A*^*c*.2299delG/c.2299delG^). **A)** Gene expression analysis indicates successful differentiation towards PPCs. The decrease in *NANOG* expression is indicative for loss of pluripotency, while the increased expression of photoreceptor markers *CRX*, *NRL*, *OPN1SW*, *OPN1LW* and *RHO* are indicative for the successful differentiation towards photoreceptor cells. **B)** RT-PCR analysis of *USH2A* exons 11 to 15 in untreated PPCs of a patient homozygous for the c.2299delG mutation in *USH2A* only revealed an amplicon containing exons 11 to 15. Continuous treatment of PPCs with QR-421a (28 days) specifically induced the skipping of *USH2A* exon 13 from the transcript. There was no evidence of partial exon 13 skipping, or the skipping of multiple exons. **C)** Quantitative analysis of *USH2A* exon 13 skipping upon continuous treatment with QR-421a. Treatment was started at 3 months post differentiations, and lasted for 28 days. Half of the culture medium was refreshed every other day with medium containing QR-421a. Skipping of exon 13 was already observed at the lowest concentration, and further increased with increasing concentrations. Data is shown as mean ± SD of 3-4 biological replicates per condition. Asterisks indicate significant differences with scrambled control oligo treated cells (* P<0.05; *** P < 0.001; **** P<0.0001; One-Way ANOVA followed by Sidak’s Multiple Comparison Test)

Next, patient-derived PPCs were treated with a continuous concentration of QR-421a for 28 days upon gymnotic delivery. Every 2 days, half of the culture medium was replaced by fresh medium containing a novel dose of QR-421a. Untreated PPCs and PPCs treated with a control oligonucleotide (with the same chemistry and length but a random sequence) were used as negative controls. RT-PCR analysis of *USH2A* exons 11 to 15 revealed that, in contrast to previous analysis in patient-derived fibroblasts (Lenassi *et al*, 2014), no alternatively spliced *USH2A* transcripts could be detected in untreated PPCs homozygous for the c.2299delG mutation (**Fig. 4B)**. Results furthermore showed that QR-421a induced significant levels of exon 13 skipping at all concentrations tested (1-10 μM), while exons 12 and 14 were retained within the *USH2A* Δexon13 transcript (**Fig. 4B, C**). At a 1 μM concentration, exon 13 skipping was observed in 42±11 % (P=0.001, Sidak’s multiple comparison test) of *USH2A* transcripts. This increased to 63±8 % (P<0.0001, Sidak’s multiple comparison test) of transcripts lacking exon 13 when QR-421a was supplied at a 10 μM concentration (**Fig. 4C**). No exon 13 skipping was detected in untreated or scrambled control oligonucleotide-treated PPCs, indicating that skipping of this exon was specifically induced by QR-421a.

### Retinal uptake and distribution of QR-421a in cynomolgus monkeys

To assess QR-421a uptake and distribution in the primate retina, cynomolgus monkeys (*Macaca fascicularis*) received a single bilateral intravitreous (IVT) injection of a low, mid and high dose of QR-421a. The distribution half-life of QR-421a in the primate retina was determined after a single (bilateral) intravitreal delivery of dose 1 or dose 2 of QR-421a. Ocular tissues from both eyes were collected at 1 hour, 12 hours, 15 days and 102 days post injection for assessment of QR-421a concentrations (n=2 monkeys per dose per time-point). The mean ocular tissue concentration of QR-421a according to the time after administration was plotted for retina and vitreous humor (**Fig. 5B, C**). The data show that following IVT administration the concentration of QR-421a in the vitreous humor declined as a result of uptake by the surrounding ocular tissues. This was illustrated by the approximately 4-fold increase of the mean QR-421a concentration in the retina at 12 hours post injection, and simultaneous decrease of QR-421a concentration in the vitreous humor (**Fig. 5B, C**). Ocular tissue concentrations observed at day 1 after injection confirm an initial rapid uptake into the retina. After the initial distribution phase, QR-421a concentrations in the retina are 2 to 3 orders of a magnitude higher than in vitreous humor. QR-421a has a long tissue residence time, with an estimated half-life time >200 days in the retina.

**Figure 5:**
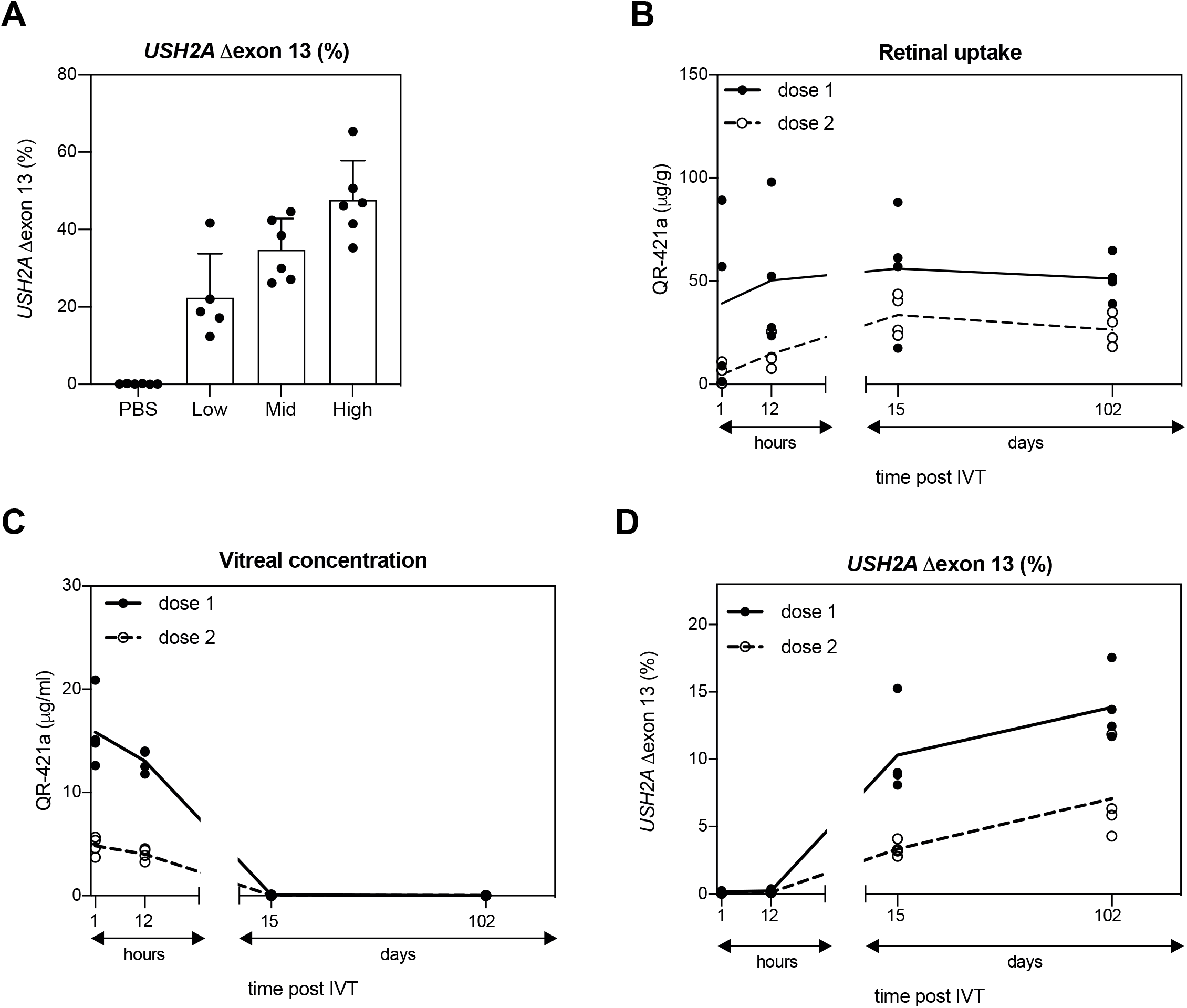
Characteristics of QR-421a upon intravitreal injection in cynomolgus monkeys. **A)** Cynomolgus monkeys received a single bilateral IVT injection of mid, low or high dose of QR-421a or PBS control and were maintained for 28 days. Retinae were collected and used to determine levels of exon 13 skipping in *USH2A* transcripts by a transcript-specific ddPCR. QR-421a induced a dose-dependent increase in *USH2A* exon 13 skipping. Data is shown as mean ± SD, and as individual values. **B)** Cynomolgus monkeys received a single bilateral IVT injection of dose 1 or dose 2 of QR-421a and were maintained for 1 hour, 12 hours, 15 days and 102 days post-injection to determine retinal uptake (n=2 monkeys/sex/time point; both eyes used for tissue concentration analysis). Retinal tissue concentration observed at the first time point confirms an initial rapid uptake into the retina. **C)** The concentration of QR-421a in vitreous humor declined before the first time point. An ELISA method was used to quantify QR-421a concentrations in the retina **(B)** and vitreous humor **(C)**. Data is shown as mean and invidual values. **D)** To determine the duration of action of QR-421a, cynomolgus monkeys received a single bilateral IVT injection of dose 1 or dose 2 of QR-421a and were maintained for 1 hour, 12 hours, 15 days or 102 day. *USH2A* transcript analysis was performed using a transcript-specific ddPCR analysis. QR-421a-mediated *USH2A* exon 13 skipping was maintained for at least 102 days. Data is shown as mean and invidual values. Abbreviations: IVT: intravitreal injection; PBS: phosphate buffered saline.

The *USH2A* gene is well conserved across species with a very high sequence similarity between humans and non-human primates. In fact, the QR-421a hybridization sequence in human and cynomolgus monkey *USH2A* is identical. In addition, the retinal anatomy of human and cynomolgus monkey is highly comparable (Slijkerman *et al*, 2015). Therefore the *in vivo* dose-dependent efficacy of QR-421a was assessed in cynomolgus monkeys which received single bilateral IVT injections of one of three dose levels of QR-421a, and exon 13 skipping was assessed at 29- and 92-days post-injection. QR-421a induced a dose-dependent *USH2A* exon 13 skipping (**Fig. 5A**). At 29 days post-injection, exon 13 skipping was detected in 26%, 48% and 69% of *USH2A* transcripts at the low, mid, and high doses, respectively. At 92 days after treatment with high dose of QR-421a, the exon skipping percentage was 79%. No exon skipping (<0.5 %) was detected in PBS treated animals. A more conservative estimate of QR-421a induced *USH2A* Δexon 13 skipping, calculated using exon 13 transcript levels from PBS treated control animals, resulted in 22%, 35% or 48% of skipping at low, mid, or high doses, respectively. Time-response of QR-421a mediated exon 13 skipping in the retina of cynomolgus monkeys was also determined. Cynomolgus monkeys received a single bilateral IVT injection at doses of dose 1 and dose 2 of QR-421a. Retinae were collected at 1 hour, 12 hours, 15 days, and 102 days post-injection for the assessment of *USH2A* exon 13 skipping levels. No exon skipping (<0.5 %) was noticed in the cynomolgus monkey retina at 1 hour and 12 hours post-injection in either of the dosed groups (**Fig. 5D**). In animals treated with dose 2 of QR-421a, 3% exon 13 skipping was detected after 15 days, which increased to 7% at 102 days. Monkeys treated with dose 1 of QR-421a showed 10% exon 13 skipping after 15 days, which increased to 14% at 102 days, after which levels of exon skipping slowly declined. These data provide *in vivo* evidence for a QR-421a-mediated dose-dependent *USH2A* exon skipping.

## Discussion

Mutations in exon 13 of the *USH2A* gene, including the founder mutations c.2299delG and c.2276G>T, are estimated to underlie syndromic (Usher syndrome) and non-syndromic retinitis pigmentosa (RP) in approximately 16,000 individuals in the Western world. In this study we used PMOs targeting *ush2a* exon 13 to evaluate exon skipping as a therapeu tic strategy for the future treatment of *USH2A*-associated RP. We show that skipping of *ush2a* exon 13 resulted in a partially restored expression of usherin in photoreceptors of *ush2a^rmc1^* larvae. Furthermore, exon 13-skipping in zebrafish resulted in electroretinogram (ERG) b-wave traces comparable to wildtype larvae, which indicates an improved retinal function. We established QR-421a, an antisense oligonucleotide drug that induces skipping of human *USH2A* exon 13 in cell and animal models, and shows a long residence time in the primate retina after a single intravitreal delivery. Our study therefore provides proof of concept for exon skipping as a highly promising treatment option for *USH2A*-associated retinal degeneration as a consequence of mutations in exon 13.

In the inner ear, usherin is essential for the maturation of hair bundles that are located at the apex of hair cells (Michalski *et al*, 2007). In the retina, the large extracellular tails of usherin and ADGRV1 have been proposed to interact and together bridge the gap between the opposing membranes of the photoreceptor connecting cilium and the periciliary region (Maerker *et al*, 2008; Sorusch *et al*, 2017). In contrast to the situation in the inner ear hair cells, usherin seems redundant for the initial development of photoreceptors (Maerker *et al*, 2008), and rather fulfills a post-developmental role. As such, usherin seems to be particularly important for photoreceptor maintenance (Liu *et al*, 2007). Therefore, therapeutic strategies that rescue the expression of functional usherin protein in the retina, can potentially prevent or slow down the progression of photoreceptor degeneration and, as such, preserve visual function in patients.

The in-frame skipping of exons harboring pathogenic variants has already been shown to have a particularly high therapeutic potential for large genes encoding (structural) proteins that contain series of repetitive protein domains (Niks & Aartsma-Rus, 2017). Duchenne Muscular Dystrophy (DMD) is caused by mutations in the *DMD* gene. *DMD* encodes dystrophin, a structural linker protein consisting of a stretch of 24 spectrin-like domains flanked by protein-protein interaction domains that are used to connect the F-actin cytoskeleton to β-dystroglycan. Exon skipping was previously shown to restore the reading frame in patients suffering from DMD due to mutations in exon 51, and has the ability to restore the production of a functional dystrophin protein (Mendell *et al*, 2016; 2013). Like dystrophin, usherin is also a large structural protein and contains repetitive EGF-lam and FN3 domains. The in-frame exon 13 of *USH2A*, containing the recurring mutations c.2299delG and c.2276G>T, encodes multiple EGF-lam domains that are proposed to form a stiff rodlike element (Beck *et al*, 1990; Yurchenco & Cheng, 1993). *In silico* predictions and 3D homology modelling demonstrated that EGF-lam domains 4 and 8, that are both in part encoded by exon 13, can fuse into a properly folded EGF-like domain after skipping of *USH2A* exon 13. The spacing between cysteine residues 5 and 6 in the resulting EGF-like 4-8 fusion domain of usherinΔexon13 is 16 amino acids, which differs from the reported consensus spacing (8 amino acids) in EGF-like domains (Appella *et al*, 1988). However, a certain variation in spacing between the fifth and sixth cysteine residue in EGF-like domains is tolerated and has been reported. For example, the spacing between cysteine residues 5 and 6 in EGF-like domains 1 and 2 of the transmembrane protein UMODL1 is 18 and 15 amino acids, respectively (Nishizaki *et al*, 2009).

Functionality of the usherinΔexon13 protein in the retina was assessed after the skipping of exon 13 in our recently published *ush2a^rmc1^* zebrafish mutant, which has a homozygous protein truncating mutation in exon 13 (Dona *et al*, 2018). Microinjection of a combination of two PMO-based antisense oligonucleotides in the yolk of single-cell staged zebrafish embryos induced a transient skipping of exon 13 from the *ush2a* transcript. As a result, a small but significant increase in usherin labeling at the photoreceptor periciliary membrane of PMO-treated *ush2a^rmc1^* larvae was observed. This indicates that translation of *ush2a* Δexon13 transcripts results in a shortened usherin protein that is able to properly localize in zebrafish photoreceptor cells. Furthermore, PMO-induced skipping of the mutated exon 13 resulted in completely restored ERG b-wave amplitudes, indicative of a restored visual function. Although PMOs are remarkably stable in zebrafish embryos, the intranuclear PMO concentration declines with the increasing number of nuclei during development (Eisen & Smith, 2008). As such, the effect of PMO-induced gene knockdown or splice modulation at the transcriptional level are generally most prominent within the first 3 days of zebrafish development (Bill *et al*, 2009). The observed levels of exon 13 skipping (^~^20% at 3dpf) already indicate that relatively few *ush2a* Δexon13 transcripts are required to rescue the retinal defects seen in *ush2a^rmc1^* larvae. As it is only useful to record ERGs in zebrafish larvae that have a functional retina (≥ 5dpf) (Biehlmaier *et al*, 2003), the complete restoration of ERG defects observed in PMO-treated *ush2a^rmc1^* larvae either suggests that even lower levels of *ush2a* Δexon13 transcripts are sufficient for retinal function, or that the encoded usherinΔexon13 is relatively stable, at least until 5-6dpf. Rods do not significantly contribute to the zebrafish ERG until 15 dpf and therefore all responses recorded in these larvae are expected to be cone-derived (Branchek, 1984; Bilotta *et al*, 2001). Patients with *USH2A*-associated RP often present with night-blindness as the initial symptom of retinal dysfunction, indicating a primary dysfunction of the rods. However, it was recently reported that both rod and cone responses were markedly reduced in the ERGs of adolescent USH2a patients (Sengillo *et al*, 2017). Therefore, a restored ERG response in zebrafish *ush2a^rmc1^* larvae upon exon 13 skipping is promising for a beneficial effect in patients. Furthermore, the functionality of the usherinΔexon13 protein is also supported by a recent book chapter by Pendse et al, demonstrating that auditory function is not affected in *Ush2a*^Δexon13/Δexon13^ nor in *Ush2a*^Δexon13/mut^ mice (Pendse *et al*, 2019).

Following the therapeutic proof-of-concept obtained for *ush2a* exon 13 skipping in zebrafish photoreceptors, we present evidence for the pharmacodynamic potential of QR-421a, the lead candidate AON for the future treatment of patients with RP due to mutations in exon 13 of the *USH2A* gene. QR-421a treatment resulted in a concentration-dependent *USH2A* exon 13 skipping in a retinoblastoma cell line. In general, retinal tissue displays a high degree of transcriptional activity and alternative splicing (Farkas *et al*, 2013). Retinal organoids and photoreceptor progenitor cells differentiated from patient-derived induced pluripotent stem cells (iPSCs) provide an excellent platform to test therapeutic interventions for inherited retinal dystrophies *in vitro*, as recently demonstrated by us and others (McDougald *et al*, 2016; Dulla *et al*, 2018). Treatment of PPCs derived from a patient homozygous for the *USH2A* c.2299delG mutation with QR-421a resulted in a dosedependent skipping of exon 13, without inducing unwanted alternative exon skipping events.

It is important that oligonucleotides intended for the treatment of retinal dystrophies are capable of accessing retinal cells in order to reach the intended target site. In this study, ocular pharmacokinetics of QR-421a administered by IVT injection were characterized in the cynomolgus monkey. The eyes of this non-human primate are highly similar to those of human in terms of retinal architecture and barriers for ocular drug delivery such as the internal limiting membrane that separates the retina from the vitreous body (Slijkerman *et al*, 2015; Peynshaert *et al*, 2019). Upon intravitreal (IVT) delivery, QR-421a is rapidly distributed into the retina. The ocular pharmacokinetics of QR-421a are similar to those described for other large hydrophilic molecules, with the vitreous acting as a central compartment from which the oligonucleotide is rapidly taken up by the surrounding ocular tissue after IVT injection, and from which it is slowly cleared via anterior and/or posterior routes (Del Amo *et al*, 2017). QR-110, an antisense oligonucleotide designed for the *CEP290*-associated splice correction therapy to treat Leber’s Congenital Amaurosis (LCA), is similar to QR-421a in chemical composition, observed exon-skipping efficiencies in human photoreceptor progenitor cells and uptake into cynomolgus monkey photoreceptors (Dulla *et al*, 2018). As an intravitreal injection of QR-110 was recently shown to improve light-perception in LCA10 patients (Cideciyan *et al*, 2019), our data indicate that QR-421a has the appropriate pharmacological properties for future clinical applications.

A maximum percentage of ~20% of *ush2a* Δexon13 transcripts was observed in PMO-treated *ush2a^rmc1^* zebrafish larvae relative to the total amount of *ush2a* transcripts in untreated strain- and age-matched wildtype larvae. As this relatively low percentage of exon 13 skipping still resulted a complete restoration of ERG traces, it is tempting to speculate on the minimal amount of *USH2A* Δexon13 transcripts needed for a detectable and durable therapeutic effect. Individuals that carry a heterozygous loss of function mutation in *USH2A* are asymptomatic, indicating that about 50% of usherin could theoretically be enough for a sustained retinal function. The work of Pendse *et al* shows that the amount of protein produced from a single *Ush2a* Δexon13 allele is indeed sufficient for normal usherin localization in photoreceptors, normal hair cell development and normal auditory function in mice (Pendse *et al*, 2019). The surprisingly low amount of usherinΔexon13 protein detected in the retina of PMO-injected *ush2a^rmc1^* larvae, nevertheless leading to complete restoration of retinal function, suggests that an even lower amount of usherinΔexon13 protein can be sufficient for retinal function. Quantitative RT-PCR analysis revealed that the amount of *ush2a* Δexon13 transcripts in PMO-treated larvae at 3dpf comprises approximately 20% of total *ush2a* transcripts in wildtype larvae. Interestingly, studies in a humanized mouse model for USH1c showed that ~20% of correctly-spliced *Ush1c* transcripts, observed after the delivery of splice-correcting AONs at postnatal day 5, is sufficient to rescue auditory function up to 3 months post injection (Lentz *et al*, 2013). Based on this, and what is known from other AONs acting through an exon skipping mechanism (Rigo *et al*, 2014; Charleston *et al*, 2018), exon skipping levels in the range of 10-20% could potentially be enough to result in sufficient protein restoration to reach efficacious levels.

The age of onset and slow rate of progression of *USH2A*-associated RP leave ample opportunity for therapeutic intervention to halt of slow disease progression. All evidence indicates that *ush2a* transcripts lacking exon 13 encode a usherinΔexon13 protein with sufficient residual function to rescue visual function. QR-421a was validated in-vitro an in-vivo for the treatment of retinitis pigmentosa resulting from mutations in exon 13 of the *USH2A* gene. Translating the results obtained in cynomolgus monkey to the doses of QR-421a approved for the currently ongoing phase I/II clinical trial (NCT03780257), it is anticipated that therapeutic levels of exon skipping can be obtained in patients upon a single IVT delivery of QR-421a.

## Supplemental information

Supplemental Information includes three figures and two tables.

## Author contributions

Conceptualization: H.D., E.V., P.A., E.W.; Writing: H.D., R.S., E.V., E.W.; Investigation and Analyses: zebrafish experiments, R.S., M.D., J.Z., S.B., T.P., E.V.; 3D modeling, H.V.; Cell culture and PPC experiments, S.A., R.P.; Supervision, H.D., S.N,. H.K., P.A., E.W; Funding Acquisition, E.W.; Review, Editing & Approval of Manuscript, all authors.

## Acknowledgements

We are grateful to the patients for donating tissue for this study. We would like acknowledge that Radboud University Zebrafish Platform, and in particularly Tom Spanings and Antoon van der Horst for their excellent zebrafish husbandry. This study was financially supported by ProQR Therapeutics, the Foundation Fighting Blindness USA (grant PPA-0517-0717-RAD to E.W.), the Dutch Organisation for Scientific Research (Veni grant 016.136.091 to E.W.), the Gelderse Blinden Stichting, Stichting Ushersyndroom, and Stichting Klavertje2.

## Materials and methods

### Animals

*ush2a^rmc1^* (c.2337_2342delinsAC; p.Cys780GlnfsTer32; Dona *et al*, 2018) and strain-matched wild-type Tüpfel long fin zebrafish were bred and raised under standard conditions (Westerfield, 2000). Both adult and larval zebrafish were kept at a light-dark regime of 14 hours light: 10 hours darkness. All experiments were carried out in accordance with European guidelines on animal experiments (2010/63/EU). Zebrafish eggs were obtained from natural spawning and reared at 28.5°C in E3 embryo medium (5 mM NaCl, 0.17 mM KCl, 0.33 mM CaCl2, and 0.33 mM MgSO4), supplemented with 0.1% methylene blue. Non-human primate studies were outsourced to Covance Laboratories, USA, who obtained all relevant approvals and the studies were conducted in accordance to all regulatory standards.

### PMO design and microinjection

Phosphorodiamidate morpholino oligonucleotides (PMOs) were designed by first assessing the target sequence for SRSF2 (SC35) exonic splice enhancer sites (threshold of 3.0) using the online ESE finder 3.0 tool (Smith *et al*, 2006). Zebrafish *ush2a* exon 13-targeting PMOs were synthesized by Gene Tools, LCC (USA). PMOs were dissolved in ultrapure water at a stock concentration of 50μg/μl and stored at −20°C. One nanoliter containing 0.5 - 4pg per PMO and 0.25% (v/v) phenol red was injected into the yolk of 1- to 2-cell-stage embryos with a Pneumatic PicoPump pv280 (World Precision Instruments). After injection, embryos were raised at 28.5°C in E3 embryo medium until analysis. PMO sequences are provided in **table S1**.

### Zebrafish *ush2a* transcript analysis

Total RNA was isolated from pools of 10-15 larvae per condition. Larvae were snap-frozen on liquid nitrogen and subsequently homogenized in Qiazol reagent (Qiagen, #79306) using a 25G needle. Upon Qiazol extraction of total RNA was performed conform manufacturers’ instruction. Total RNA was further purified and DNase treated using the Nucleospin RNA extraction kit (Machery-Nagel, #740955.50). One microgram of total RNA was reverse transcribed using SuperScript VILO reverse transcriptase kit (ThermoFisher Scientific, #11755050). PMO-induced alternative splicing of *ush2a* transcripts was analysed by PCR amplification using primers in zebrafish *ush2a* exon 11 using Q5 HF DNA polymerase (New England Biolabs, #M0491L). Primer sequences are provided in **table S2**. Amplified transcripts were visualized by agarose gel electrophoresis (1% agarose in 0.5xTBE) and subsequently validated using Sanger sequencing. Exon 13 skipping levels were determined using a quantitative RT-PCR approach, including a standard curve of custom synthetic oligonucleotide templates (gBlocks; Integrated DNA Technologies), using primer pairs that specifically amplify exon 13-containing *ush2a* transcripts or Δexon13 *ush2a* transcripts. Targets were amplificated using GoTaq DNA polymerase (Promega, #M3001) and analyzed on a QuantStudio 3 Real-Time PCR System (Applied Biosystems).

### Zebrafish immunohistochemistry and quantification of fluorescent signal intensity

Per group, 10 zebrafish larvae were imbedded in Tissue-Tek OCT compound (Sakura, #4583) without prior fixation. Unfixed cryosections were permeabilized using 0.01% Tween20 (Merck, #8.22184) in PBS, rinsed and then pre-incubated with blocking solution (10% Normal Goat Serum (Brunschwig, #G-S-1000) and 2% Bovine Serum Albumin (Sigma-Aldrich, #A7906) in PBS). Primary antibodies (rabbit anti-C-terminal zebrafish usherin (1:1000; Novus Biologicals, #27640002), mouse anti-centrin (1:500; Millipore, #04-1624)) were diluted in blocking solution and incubated overnight at 4°C. After rinsing the sections three times with PBS, they were incubated for 1 hour at room temperature with blocking solution containing the secondary antibodies (Alexa fluor 568 Goat anti-rabbit (1:800; Molecular Probes, #A11011) Alexa Fluor 488 goat anti-mouse (1:800; Molecular Probes, #A11029)) and the nuclei staining DAPI (diluted 1:8000; Molecular Probes, #D1306). After a dip in ultrapure water, the sections were coverslipped with Prolong Gold antifade reagent (Life Technologies, #P39930; lot.#1737358). The sections were examined using a Zeiss AxioImager Z2 microscope with ZEN 2012 software and photographed with a Zeiss AxioCam 506 mono camera.

Images of the middle section of each eye, taken at identical exposure settings, were used for quantification of the fluorescent signal intensity of anti-usherin immunoreactivity using the Fiji v1.47 software (Schindelin *et al*, 2012). First, the area of the connecting cilia was selected and manually isolated from the picture based on the centrin immunofluorescence signal. Subsequently, a mask was made based on the centrin staining using the ‘Find Maxima’ option (noise□=□50), and dilated five times. To find the exact location of usherin immunofluorescence, the centrin mask and layer containing the usherin immunofluorescent signal were combined. Find Maxima (noise□=□10) was used to identify the usherin immunofluorescence within the centrin mask. The resulting mask was dilated three times and touching objects were separated using the watershed option. Subsequently, the maximum gray value of the identified regions was measured on the original image of usherin immunofluorescence (“Analyse Particles” option; size□=□0–50, pixel circularity□=□0.00–1.00).

### Electroretinogram recordings in zebrafish larvae

Larvae, of 5-6 days post-fertilization (dpf), were placed on a filter paper in the middle of a plastic recording chamber. The chamber contained 1% agarose, in which the reference electrode was inserted. The isolated eye was positioned to face the light source. Under visual control via a standard microscope equipped with red illumination (Stemi 2000C, Zeiss, Oberkochen, Germany), a glass microelectrode with an opening of approximately 20 μm at the tip was placed against the center of the cornea. This electrode was filled with E3 medium (5 mM NaCl, 0.17 mM KCl, 0.33 mM CaCl, and 0.33 mM MgSO4), the same in which the embryos were raised and held. A custom-made stimulator was invoked to provide light pulses of 100 ms duration, with a light intensity of 7000 lux. It uses a ZEISS XBO 75W light source and a fast shutter (Uni-Blitz Model D122, Vincent Associates, Rochester, NY, USA) driven by a delay unit interfaced to the main ERG recording setup. Electronic signals were amplified 1000 times by a pre-amplifier (P55 A.C. Pre-amplifier, Astro-Med. Inc, Grass Technology) with a band pass between 0.1 and 100 Hz, digitized by DAQ Board NI PCI-6035E (National Instruments) via NI BNC-2090 accessories and displayed via a self-developed NI Labview program.

### AON delivery in a retinoblastoma cell line

The WERI-Rb1 (ATCC^®^ HTB-169™) retinoblastoma cell line was obtained from ATCC. WERI-Rb1 cells were cultured in RPMI 1640 medium (Gibco, #21875034) supplemented with 10% fetal bovine serum (Bio-West, #S1810-500). Cells were maintained by addition of fresh medium or replacement of medium every 3 to 4 days.

Cells were transfected with QR-421a using Lipofectamine 2000 transfection reagent (Invitrogen, #11668019). Concentrations of QR-421a used were: non-treated (only Lipofectamine 2000), 10 nM, 25 nM, 50 nM, 100 nM and 200 nM. As a control, cells were transfected with 200 nM of scrambled AON (2’-MOE modification). A ratio of 2:1 (volume/weight) between Lipofectamine 2000 and the AON was used. Both Lipofectamine 2000 and AON were prepared in Opti-MEM. Lipofectamine 2000 mixture was added to AON mixture and incubated for 20 minutes at room temperature before adding the transfection complexes to the cells. Cells were incubated for 24 hours at 37°C. Two samples were treated per condition.

For gymnotic delivery of QR-421a, concentrations of QR-421a used in WERI-Rb1 cells were: non-treated (only medium), 10 μM, 25 μM, 40 μM and 50 μM. As a control, cells were treated with 50 μM scrambled AON (2’-MOE modification). AONs were prepared in the desired concentration in culture medium and added to the cells. Cells were incubated for 48 hours at 37°C. Two samples were treated per condition.

### Differentiation and AON treatment of iPSC-derived photoreceptor progenitor cells

Fibroblasts were reprogrammed using 4 lentiviruses expressing *Oct3/4*, *Sox2*, *Klf4* and *c-Myc*. iPSC lines were generated on feeder cells (mouse embryonic fibroblasts), and subsequently maintained in Essential 8 medium (Life Technologies, #A1517001). After reaching confluence, iPSC clumps were digested with accutase (Sigma-Aldrich, #A6964) and plated in a 12-well plate to form a monolayer. Upon reaching confluence Essential-Flex E8 medium (Thermo Fisher Scientific, #A2858501) was changed into a differentiation medium consisting of DMEM/F12 (Gibco, #11320-033), supplemented with non-essential amino acids (NEAA; Gibco, #11140-050), B27 supplements (Thermo Fisher Scientific, #1287010), N2 supplements (Thermo Fisher Scientific, #17502048), 100 ng/μL insulin-like growth factor-1 (IGF-1; Sigma-Aldrich, #I3769), 10 ng/μL recombinant fibroblast growth factor basic (βFGF; Sigma-Aldrich, #F0291), 10 μg/μL Heparin (Sigma-Aldrich, #H3149-10KU) and 200 μg/mL human recombinant COCO (Bio-Techne, #3047-CC). The medium was changed every day for 90 days, after which the cells were treated with different concentrations of AONs during 1 month. At the end of the 4^th^ month the cells were collected and characterized.

Photoreceptor precursor cells (PPCs) were treated continuously with 1, 2, 5 or 10 μM QR-421a or 10 μM control oligo (which has the same chemistry and length but a random sequence). Treatment was started after 90 days of differentiation and lasted for 28 days. Every two days, 50% of culture medium was refreshed with fresh culture medium containing AON.

Quantitative RT-PCR was used to evaluate the differentiation status at the end of the experiment. Expression of *NANOG*, *CRX*, *NRL*, *OPN1SW*, *OPN1LW* and *RHO* was investigated using GoTaq DNA polymerase (Promega, #M3001) and a QuantStudio 3 Real-Time PCR System (Applied Biosystems). Data are normalized for the expression of the housekeeping gene *GUSB*. Primer sequences are provided in **table S2**.

### Human *USH2A* transcript analysis

Total RNA was isolated from the cells using RNeasy Plus Mini Kit (Qiagen, #74136) according to manufacturer’s protocol and cDNA was synthesized. The visualize AON-induced alternative splicing in WERI-Rb1 cells and human iPSC-derived photoreceptor progenitor cells, a PCR was performed using primers on *USH2A* exons 11 and 15. PCR fragments were gel-extracted, purified using the NucleoSpin^®^ Gel and PCR Clean-up kit (Machery-Nagel, #740609.50) and subjected to Sanger Sequencing.

For the quantification of *USH2A* transcripts, digital droplet PCR (ddPCR) was performed using the One-step RT-ddPCR advanced kit for probes (Bio-Rad, #1864022). The final 20 μL reaction mix contained: 5 μL Supermix, 2 μL Reverse Transcriptase, 1 μL 300 mM DTT, 1x TaqMan gene expression assays or 450 nM *USH2A* forward and reverse primer each and 250 nM *USH2A* probe. Total *USH2A* (VIC label) and *USH2A* exon 13 (FAM label) levels were quantified using 50 or 100 ng RNA in a multiplex manner using commercial TaqMan gene expression assays. *USH2A* Δexon 13 levels were quantified in 50 or 100 ng RNA using an inhouse designed assay. PCR reactions were dispersed into droplets using the QX200 droplet generator (Bio-Rad) according to the manufacturer’s instructions and transferred to a 96-well PCR plate. End point PCR was performed in a T100 Thermocycler (Bio-Rad). The fluorescence of each droplet was quantified in the QX200 droplet reader (Bio-Rad). Each sample was analyzed in duplicate. Absolute quantification was performed in QuantaSoft software (Bio-Rad). Thresholds were manually set to distinguish between positive and negative droplets.

For data analysis, average copy numbers of duplicate measurements per ng RNA input were calculated and the target copy number was normalized for total *USH2A*. For this, the target gene copy numbers were divided by the total *USH2A* copy numbers and multiplied by the average total *USH2A* copy number in that run. Percentage of *USH2A* Δexon 13 transcripts was calculated using the following scheme: Treated sample’s Δexon 13 transcripts divided by average sum of untreated samples’ exon 13 and Δexon 13 transcripts, multiplied by 100. In case untreated samples were not available, control treated samples were used. This method was used as multiple *USH2A* isoforms are present in the cells and the splicing modulation may not only lead to exon 13 skipping (*e.g*. combined exon 12 and 13 skipping, partial exon 13 skipping).

### QR-421a quantitation in tissue samples by ELISA

The PK of QR-421a was evaluated in cynomolgus monkeys after IVT administration. A dual hybridization ELISA method (hELISA) was used to quantify QR-421a concentration in ocular tissues. The hELISA method is a 2-step enzyme linked immunosorbent (ELISA) procedure and runs on the MSD ECL platform with commercially available reagents. Detection is performed by measuring light emitted by a sulfo-tag labeled antibody. The methods were demonstrated to be linear within a relevant concentration range and quantification limits were relevant to the characterization of the QR-421a tissue and serum concentration versus time profiles.

---

**Supplemental figure 1:**
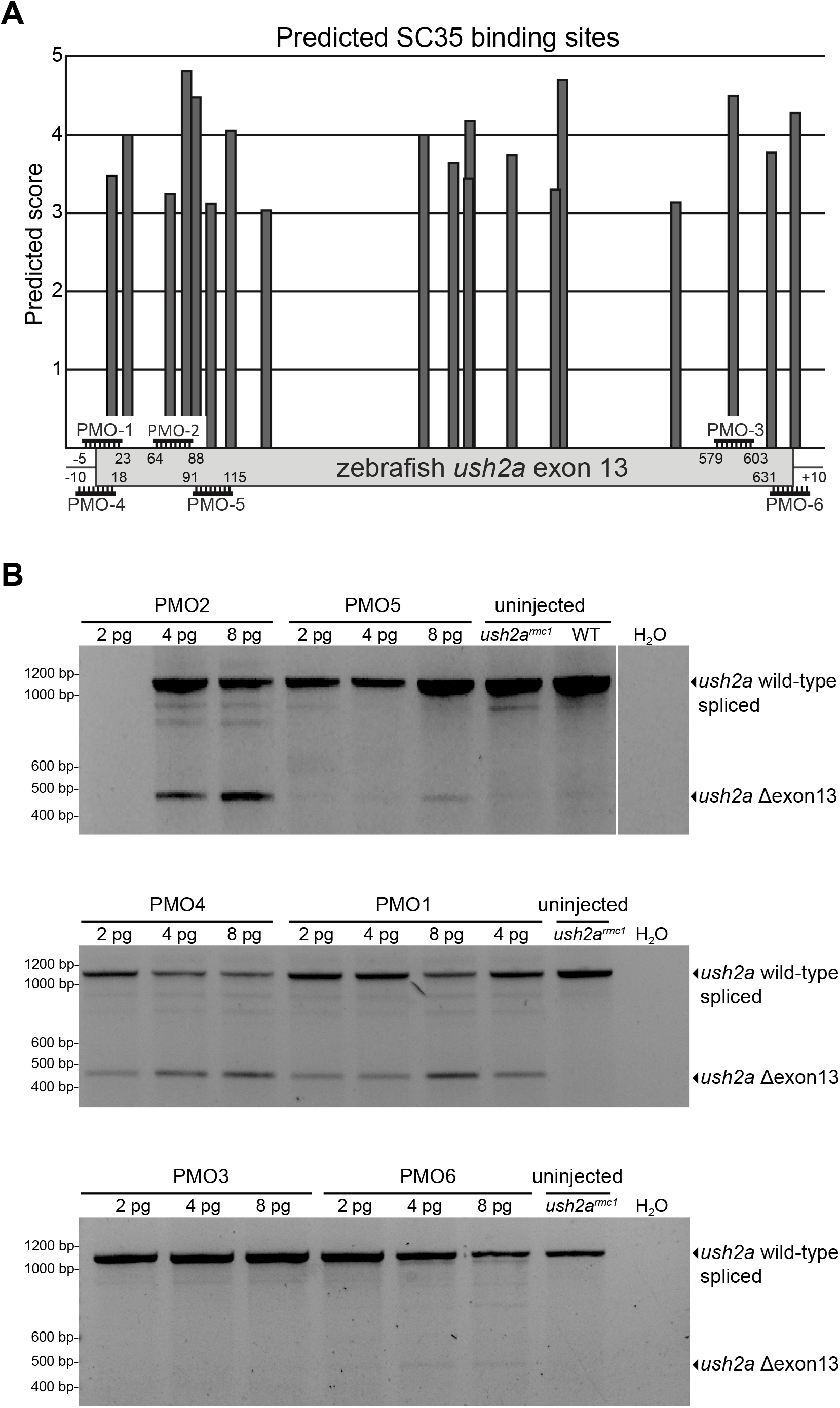
Design and validation of PMOs to induce skipping of *ush2a* exon 13 in zebrafish. **(A)** SC35 splice factor binding sites in ush2a exon 13. Target sites of the individual PMOs are indicated in the plot. Numbers indicate the first and last nucleotide of the PMO target sequences in zebrafish *ush2a* exon 13. The bars represent the presence and predicted strength of SC35 splice factor binding sites. Prediction score are indicated on the Y-axis and the ush2a sequence on the X-axis. (B) Test of the exon-skipping potential of individually delivered PMOs in zebrafish *ush2a^rmc1^* mutant larvae. All PMOs were injected in the yolk of 1- to 2-cell-stage zebrafish embryos, and investigated for exon 13-skipping potential at 3dpf (PMO2 and PMO5) or 5dpf (PMO1, PMO3, PMO4, PMO6) using RT-PCR.

**Supplemental figure 2:**
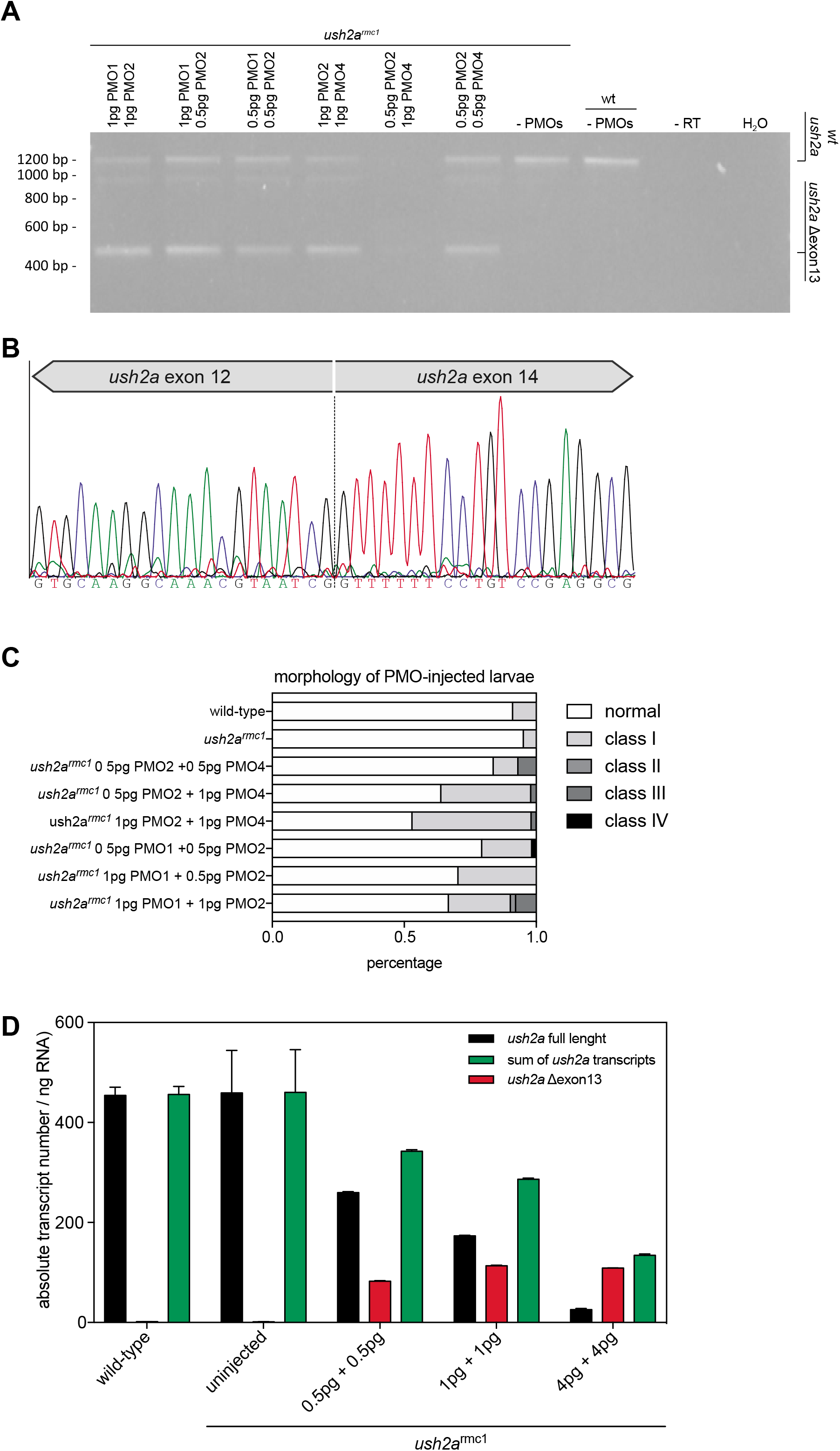
identification of the most efficient PMO or PMO pair to induce skipping of *ush2a* exon 13 in zebrafish larvae. **A)** Analysis of the exon-skipping potential of two PMO combination in zebrafish *ush2a^rmc1^* mutant larvae. All PMOs were injected in the yolk of 1- to 2-cell-stage zebrafish embryos, and investigated for exon 13-skipping potential at 3dpf. **B)** Sanger sequencing traces of the *ush2a* Δexon13 amplicon, confirming the perfect skipping of exon 13 from the transcript. **C)** Morphological analysis of PMO injected larvae at 5dpf. Phenotypic changes are classified as follows: pericardial edema (class I), body axis curvature (class II), combination of pericardial edema and body axis curvature (class III) and severe morphological malformations (class IV). Phenotypic distribution is presented as the percentage of each phenotypic group, within a total of 50-60 larvae per condition. **D)** Quantitative analysis of wildtype *ush2a* and *ush2a* Δexon13 transcripts using absolute quantitative RT-PCR. Upon increasing doses of PMOs, the amount of *ush2a* Δ exon13 transcripts hardly increases, whereas the levels of wildtype transcripts are markedly decreased. Data are presented as mean ± SD of 2 pools of 10-15 larvae.

**Supplemental figure 3:**
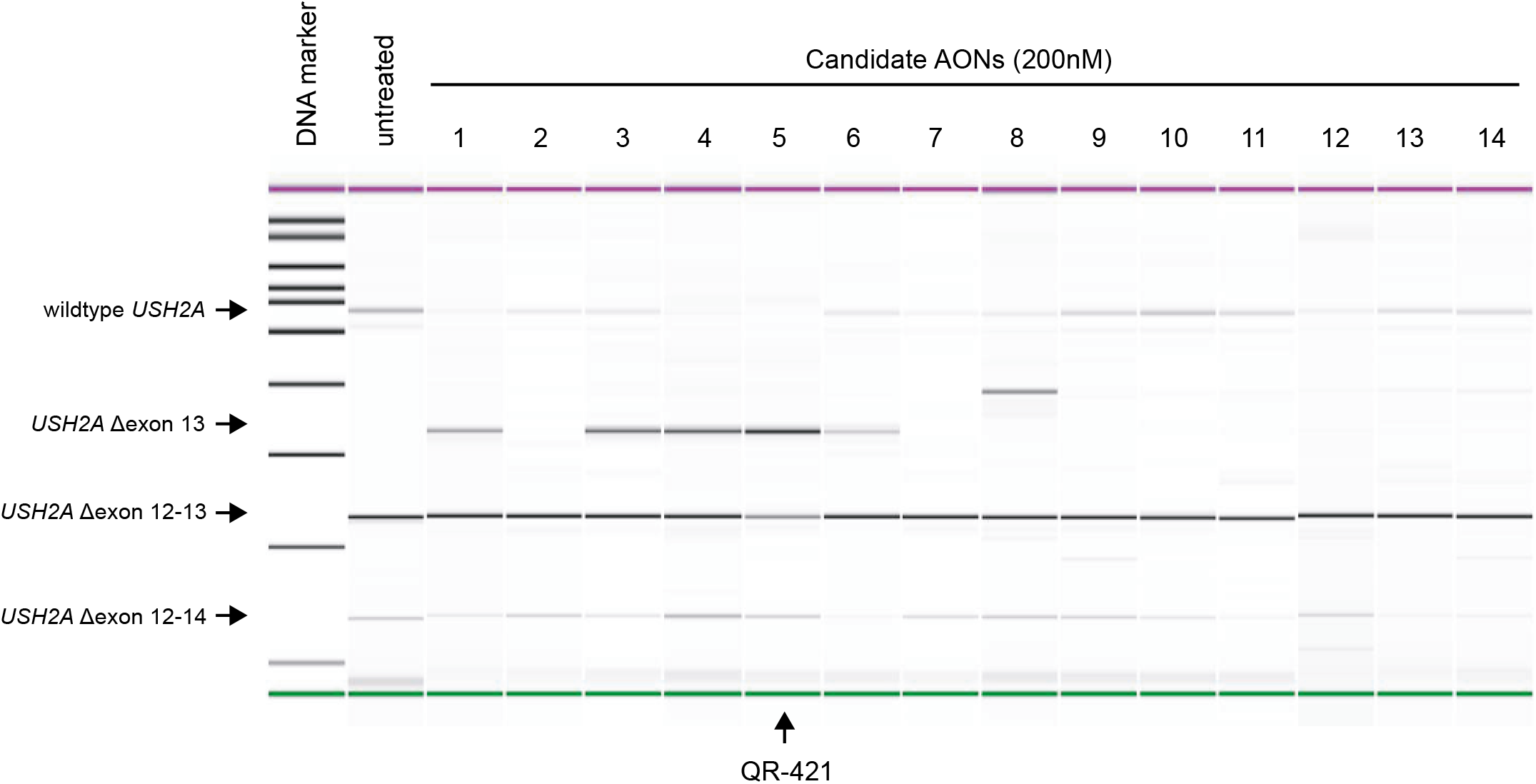
AON screening to identify the most potent AON sequence. WERI-Rb1 retinoblastoma cells were transfected with 14 different AONs (200nM), and analyzed 24 hours post transfection for their *USH2A* exon 13-skipping potential. Note that the co-skipping of *USH2A* exons 12-13 and exons 12-14 occurs naturally in WERI-Rb1 cells. AONs 1, 3, 4, 5 and 6 were able to induce the skipping of exon 13 from the *USH2A* transcript. Candidate AON 5 was selected as the most optimal sequence for further preclinical development, as it shows the strongest exon 13 skipping potential, and does not result in the increased formation of alternatively spliced *USH2A* transcripts. All tested AONs here are 21-mer RNA antisense oligonucleotides with 2’-O-methyl sugar modification and a fully phosphorothioate backbone.

