## Supplemental tables for "Antisense oligonucleotide-based treatment of retinitis pigmentosa caused by mutations in *USH2A* exon 13"

**Table S1. Antisense morpholino oligo sequences targeting zebrafish *ush2a* exon 13.**

| **Name** | **Sequence** | **Targeted motifs** |
| --- | --- | --- |
| PMO-1 | 5’-GTTACAACGGTCACAGGTTAGACCTAAA-3’ | Splice acceptor and SC35 motif 1 |
| PMO-2 | 5’-CATGGGTCACAGCCACAGGAAATGC-3’ | SC35 motif 3 and 4 |
| PMO-3 | 5’-GGTAGGCAGATACACTGACCACTTA-3’ | SC35 motif 18 |
| PMO-4 | 5’-AACGGTCACAGGTTAGACCTAAAAATAA-3’ | Splice acceptor and SC35 motif 1 |
| PMO-5 | 5’-GGATTACAGAACTGGTGCAGAGAAC-3’ | SC35 motif 5 and 6 |
| PMO-6 | 5’-AAGCACTAACCTGGTTTACAGGTTCCAC-3’ | Splice donor and SC35 motif 19 and 20 |
| Control PMO | 5’-CCTCTTACCTCAGTTACAATTTATAC-3’ | - |

**Table S2. Primer list**

| **Target** | **Species** | **Sequence** |
| --- | --- | --- |
| *ush2a* exon 11-14 | Zebrafish | 5’-AGCGCTGTCGGAGTCTCTTC-3’  5’-CTGTGACCGGTCAGTGATGG-3’ |
| *ush2a* exon 12+13 | Zebrafish | 5’-TGTATCTGCCTACCCACACG-3’  5’-CACACACACACTGCCCTGA-3’ |
| Δexon13 *ush2a* | Zebrafish | 5’-AGTGCAATCAGTGCCAACAC-3’  5’-CGGACAGGAAAAAACCGATTAC-3’ |
| *NANOG* | Human | 5’-CCTGTGATTTGTGGGCCTG-3’  5’-CAGTCTCCGTGTGAGGCAT-3’ |
| *CRX* | Human | 5’-GCCCCACTATTCTGTCAACG-3’  5’-CTTCAGAGCCACCTCCTCAC-3’ |
| *NRL* | Human | 5’-GGCTCCACACCTTACAGCTC-3’  5’-AGCCAGTACAGCTCCTCCAG-3’ |
| *OPN1SW* | Human | 5’-ACCATTGGTATTGGCGTCTC-3’  5’-GGAGAGAGGCACAATGAAGC-3’ |
| *OPN1LW* | Human | 5’-GTGGTCACTGCATCCGTCTT-3’  5’-ACGGTCTCTGCTAGGTCAGC-3’ |
| *RHO* | Human | 5’-TCATCATGGTCATCGCTTTC-3’  5’-CATGAAGATGGGACCGAAGT-3’ |
| *GUSB* | Human | 5’-TGTTTCGGTTGGTTGCCTCC-3’  5’-GGTCCAGGTTTGTCCTCTGC-3’ |
| *USH2A* exon 11-15 | Human | 5’-AGTTGGTGCAGATCCTTCGG-3’  5’-CTTGCACTGGGAACACAAGC-3’ |
